# Evaluation of Dorado v5.2.0 *de novo* basecalling models for the detection of tRNA modifications using RNA004 chemistry

**DOI:** 10.64898/2025.12.09.693013

**Authors:** Bhargesh Indravadan Patel, Franziskus N. M. Rübsam, Yu Sun, Ann E. Ehrenhofer-Murray

## Abstract

Direct RNA sequencing with Oxford Nanopore Technologies (ONT) captures nucleotide-specific current signals that reflect both sequence and chemical modifications, offering the potential to detect RNA modifications directly from native RNA molecules. To interpret such signals, ONT provides modification-aware basecalling models that estimate the probability of selected modifications at each nucleotide. In May 2025, ONT released updated modification-calling models (Dorado v5.2.0) for pseudouridine (Ψ), inosine, m^6^A and m^5^C, alongside new models for 2′O-ribose-methylations, necessitating independent validation. Here, we benchmark Dorado v5.2.0 against v5.1.0 using *ex cellulo* tRNAs from *Schizosaccharomyces pombe*, leveraging their well-defined modification landscape. We generated modification probability profiles at single-nucleotide resolution and quantified model performance using curated sets of annotated and validated modification sites. Our results reveal that, despite notable improvements in Ψ detection, most modification callers remain challenged by the dense and heterogeneous modification environments of tRNAs. This work provides the first comprehensive evaluation of Dorado v5.2.0 on native tRNAs and establishes a methodological framework for benchmarking future ONT modification models in complex RNA modification contexts.

## Background

Chemical modifications of RNA, collectively referred to as the “epitranscriptome”, play a crucial role in ensuring the proper function of diverse classes of RNA. While modifications on ribosomal RNAs (rRNAs) [1] and transfer RNAs (tRNAs) [2, 3] have been known for decades and mapped to specific sites, the functional significance of RNA modifications in messenger RNA (mRNA) has only emerged over the past decade [2, 4]. RNA modifications variably affect the basepairing and stacking properties of nucleobases, thus impacting RNA structure, folding, stability and RNA-protein interactions [5, 6]. The involvement of tRNA modifications in human diseases like cancer, neurological disorders and modopathies highlights the relevance of this field for human health [3, 7–10].

The biological importance of RNA modifications calls for methods to reliably detect them at single-base and single-molecule resolution. Direct RNA sequencing (dRNA-seq) using membrane-embedded biological nanopores, commercially developed by Oxford Nanopore Technologies (ONT), offers a promising approach to achieve this goal. In this technique, single RNA (or DNA) strands are threaded through a biological pore, which leads to characteristic alterations of the ion current depending on the identity of the nucleobases passing through the pore [11, 12]. Subsequent interpretation of such current signal patterns using machine learning algorithms allows the inference of the RNA sequence (commonly referred to as “basecalling”) [13]. Of note, since multiple bases reside in the nanopore at a given time, the signal generated by a single nucleobase depends not only on its identity, but also on the surrounding sequence context [14]. Training of a basecaller therefore ideally requires input signals of a particular nucleobase in any possible k-mer sequence context (k = 6 to 11) [15].

Since nanopore signals depend on the chemical properties of the nucleobase, nanopore sequencing in principle should allow the detection not only of canonical bases (G/ A/ C/ U), but also of base modifications. Two principal methods have been developed for this purpose: the *sample comparison* method and *de novo prediction* method. In the sample comparison method, signal deviations generated by base modifications compared to an unmodified control can be assessed directly [16–18]. Alternatively, systematic basecalling errors generated by the modified bases using the standard basecallers are compared to those of the unmodified control to capture modified sites [19]. In *de novo* modification basecalling, the basecallers directly call the modified nucleobase, which is achieved by training models on datasets in which the modification occurs in diverse sequence contexts. Initially, the RNA basecaller from ONT (Dorado 0.1.0, SQK-RNA002 sequencing kit) was limited to calling standard nucleotides. In December 2023, ONT introduced a new RNA chemistry (SQK-RNA004) with redesigned biological pores and released *de novo* basecalling models for pseudouridine (Ψ) and N^6^-methyladenosine (m^6^A) (May 2024), as well as 5-methyl-cytosine (m^5^C) and inosine (September 2024) (Dorado v5.1.0), reflecting the biological importance of these modifications for mRNA function.

For the *de novo* prediction by Dorado, nucleobases are first assigned to a canonical base, and the modification basecaller subsequently returns a score (integer between 0 and 255) for the modification probability of that site in an individual RNA molecule. Of note, the modification basecaller samples only those sites that were assigned to the respective canonical base, but not those that were miscalled as another canonical base. This is relevant for modifications that yield high miscalling rates, because only those reads in which the site was not miscalled, will be assigned a modification score. Such scores across a large number of reads can be represented as modification probability plots. If a site is fully modified in a sample, the modification probability plot will show mostly high modification scores, whereas non-modified sites show mostly low scores.

The release of models for *de novo* basecalling of RNA modifications necessitates an independent evaluation of their performance on biological samples. In our earlier work [20], we used direct sequencing of tRNAs isolated *ex cellulo* from *Saccharomyces pombe* to assess the capabilities of the *de novo* Dorado RNA modification basecallers (Dorado v5.1.0). tRNAs constitute a good test case for this purpose, because the sites of modification as well as the modification enzymes are well known [3]. However, individual tRNAs carry 5 to 15 modified sites [21], thus complicating the interpretation of nanopore data. Our evaluation of Dorado v5.1.0 on tRNAs showed that the Ψ caller correctly predicts 63% of the annotated Ψ sites (81 of 128 sites) and furthermore revealed 17 novel sites that were validated by dRNA-seq of samples lacking individual pseudouridine synthases. However, the Ψ caller also shows high false-positive rates in that it erroneously calls dihydrouridine and complex, Elongator-dependent U34 modifications as Ψ. Furthermore, it is sensitive to modifications of other bases at neighboring positions. In such cases, the modification probability plots often show a propensity for intermediate modification scores. The Dorado v5.1.0 inosine caller showed limited performance on tRNAs, since it reliably called only one of seven known inosines in tRNAs, and the m^5^C caller gave high false-negative and false-positive scores. Finally, the Dorado v5.1.0 m^6^A caller erroneously returned elevated scores especially on N^1^-methyl-adenosine (m^1^A) sites in tRNA (m^6^A is absent from tRNAs).

Leveraging these properties of Dorado v5.1.0, we subsequently combined the sample comparison and *de novo* basecalling methods into a single framework (termed “MoDorado”) to detect tRNA modifications beyond those supported by the modification-specific models. This was achieved by comparing the distribution of prediction scores at a particular site to the distribution of an unmodified control. We thus were able to detect seven additional modifications, namely 5-carbamoylmethyluridine (ncm^5^U), 5-methoxycarbonylmethyluridine (mcm^5^U), 5-methoxycarbonylmethyl-2-thiouridine (mcm^5^s^2^U), N^7^-methylguanosine (m^7^G), queuosine, m^1^A, and N^6^-isopentenyladenosin (i^6^A), thereby generating an expanded map of these tRNA modifications in *S. pombe* [20].

Perhaps not surprisingly, an analysis of the Dorado v5.1.0 models using synthetic oligonucleotides showed high performance of the Ψ and m^6^A callers, and evaluation on mRNA samples *ex cellulo* yielded a large number of *de novo* called modification sites (Ψ, m^6^A, m^5^C, inosine) whose validity requires further investigation [22]. Another study demonstrated high recall of Ψ and m^6^A on mRNA samples, but also high false discovery rates across different sequence contexts, making modification detection especially at sites with low modification frequency unreliable [23].

In May 2025, ONT released new basecalling models for the Ψ, inosine, m^6^A and m^5^C callers and additionally generated models for 2’O-ribose-methylations (Dorado v5.2.0), thus calling for a (re)assessment of their modification detection capabilities in the context of *ex cellulo* samples. In this work, we present a comparison of the Dorado v5.2.0 compared to v5.1.0 modification caller performance as well as the 2’O-ribose methylation callers in the context of tRNA samples from *S. pombe*. This analysis showed that the Ψ caller has gained specificity and is currently the most reliable caller, while still being sensitive to neighboring modifications. The m^5^C and inosine callers show limited performance in the context of tRNAs. The Um basecalling model is sensitive to its target modification, but also erroneously detects m^5^U sites. Altogether, our study provides a comprehensive evaluation of the latest *de novo* modification calling models, highlights their strengths and limitations, and establishes a framework for benchmarking future ONT modification models in complex modification contexts such as tRNAs.

## Methods

### Data sources

dRNA-seq datasets from *in vitro*-transcribed (IVT) *S. pombe* tRNAs and tRNAs isolated *ex cellulo* from *S. pombe* (European Nucleotide Archive accession number PRJEB88080), sequenced with the ONT direct RNA sequencing kit SQK-RNA004 [20] were re-evaluated in this study. In addition, samples BSP1 (wildtype + q), BSP3 (wildtype) and BSP5 (IVT) were sequenced on a FLO-MIN004RA flow cell using a MinION Mk1C device as previously described [20], and merged with the respective earlier datasets. A description of each dataset used in this study is shown in Supplementary Table 1.

### Data processing

#### Reference curation

To generate the reference, all unique *S. pombe* mature and intron-containing tRNAs from GtRNAdb [24] were combined with all mitochondrial tRNAs from PomBase [25]. The 5′ and 3′ splint adapter sequences were then appended to each reference sequence to reproduce the final ligated library structure used for nanopore sequencing.

#### Annotation of tRNA modifications in S. pombe

The complete set of tRNA modifications annotated sites and their sources is presented in Supplementary Table 2. They represent an update compared to Rübsam et al. 2025 [20], which compiled and annotated Ψ and other known tRNA modifications based on experimental evidence in *S. pombe* or by inference from corresponding positions in *S. cerevisiae*.

#### Basecalling and modification calling with Dorado

Basecalling was carried out using the Oxford Nanopore basecaller Dorado with the models rna004_130bps_sup@v5.1.0 and rna004_130bps_sup@v5.2.0. Modification prediction was enabled using Dorado’s integrated models, which included Ψ, m^5^C, inosine, and m^6^A for version 5.1.0, and additionally four Nm models for version 5.2.0.

#### Read alignment and filtering

All *ex cellulo* sample reads were aligned to the reference using Parasail (version 2.6.0) [26] The aligned reads were subsequently filtered as follows: (1) for each read, only top-scoring alignments mapping uniquely to a single tRNA isoacceptor were retained; (2) reads with an alignment score (AS) below 95% specificity were excluded, following the threshold criteria described previously [27]; and (3) only reads spanning the full length of the tRNA were kept for downstream analyses.

*In vitro* transcribed (IVT) tRNA sample reads were aligned to the reference using BWA-MEM [28] and filtered according to the criteria described by Lucas et al. 2024 [29]. Only reads covering the full length of the tRNA were retained for downstream analyses.

#### Defining modification likelihood (ml) score and ml score threshold for assessing the performance of de novo modification callers

For each nucleotide in the reference, modification scores are assigned as discrete integers from 0 to 255, corresponding to predicted modification probabilities ranging from 0% to 100%. Predictions were retained only at positions of the relevant canonical nucleotide (e.g., Ψ predictions were retained exclusively at reference uridine positions). To retain all probability values in the output, the default reporting threshold (5%, equivalent to a score ≥12) was overridden using the `--modified-bases-threshold 0` option. Additionally, because probabilities of 0 are not included in the output files, reads lacking a reported modification probability at a given position were assigned a value of 0.

We previously defined a prediction score threshold of 80% (score = 204) to identify Ψ-modified sites using the Ψ-caller and calculated the Ψ modification fraction as the proportion of reads with scores >204 at each U position (N reads with scores >204 / N total reads) to assess the caller’s performance [20]. We also observed that non-specific modifications frequently generate a large number of reads with intermediate probabilities (20% < Ψ probability < 80%). To reduce this ambiguity, reads with intermediate probabilities were discarded, and a modification likelihood (ml) score was defined as the fraction of reads with high modification probability:

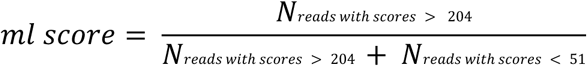

Finally, to make a binary decision on whether a reference site is modified or unmodified, the threshold for the ml score was defined as the 99th percentile of the ml score distribution derived from modification calling on IVT tRNAs.

## Results

### Dorado v5.2.0 *de novo* modification calling models for Ψ show increased sensitivity compared to v5.1.0

The release of updated models for the *de novo* basecalling of RNA modifications in Nanopore sequencing warranted an evaluation into their performance on biological samples. tRNAs can be used for this purpose, because their modification sites are well established. Nevertheless, they constitute a challenging test case owing to the high density of neighboring modifications, which complicates their detection. To directly compare the Dorado v5.2.0 modification calling model to v5.1.0, we merged independently generated datasets of tRNAs from wildtype *S. pombe* (wt + q) cultured in the presence of 100 nM queuine (queuine ensures the formation of the queuosine modification at position 34 in the relevant tRNAs [30]), and sequenced by dRNA-seq using RNA004 to achieve high coverage (total reads = 4,983,027) (Supplementary Table 2). Furthermore, as a reference for unmodified tRNAs, merged datasets of dRNA-seq from IVT *S. pombe* tRNAs were analyzed (total reads = 1,485,624). Basecalling on the two datasets (IVT, wt + q tRNAs) was performed separately using the Dorado sup v5.1.0 and v5.2.0 modification calling models.

For *de novo* prediction of RNA modifications, the Dorado basecallers first assign a canonical nucleobase to each position. The modification basecaller subsequently assigns an integer modification score (0 - 255) to the respective canonical base in an individual read, reflecting the predicted likelihood of modification at that site. For instance, the *de novo* Ψ caller only assigns a Ψ score to those sites that were basecalled as U. It was therefore of interest to determine the level of basecalling errors by the Dorado versions, because positions called as another base (e.g., G, A or C for Ψ) will not be assigned a modification score. Analysis of basecalling error rates revealed similar error frequencies between the Dorado v5.1.0 and v5.2.0 models, indicating that v5.2.0 does not substantially improve basecalling accuracy at unmodified and modified sites in tRNAs (Supplementary Fig. 1).

To assess the performance of the *de novo* callers for RNA modifications (Ψ, m^5^C, I, m^6^A), we next converted the modification probability distributions into a single modification likelihood (ml) score for each site. The ml score was defined as the ratio of the number of reads with >80% modification score divided by those with <20% modification score (see Methods). The ml score thus excludes intermediate scores, which, as we previously showed [20], often result from modifications other than the target modification (so-called “off-target” modifications).

For the Ψ caller, overall ml scores (Ψ_ml_) were high at many, though not all, known Ψ sites, with overall lower values in v5.2.0 than in v5.1.0 (Fig. 1A). At certain canonical sites such as Ψ13 and Ψ55, Ψ_ml_ scores were higher with v5.2.0, indicating improved detection in these sequence contexts (Fig. 1B, Supplementary Fig. 2). In contrast, the Ψ caller previously produced elevated scores at several non-Ψ uridine modifications – dihydrouridine (D), ncm^5^U (&, Modomics one-letter code), mcm^5^U (1), mcm^5^s^2^U (3), and Um (J) – and these off-target values were reduced in v5.2.0 (Fig. 1A). The decrease was particularly pronounced at ncm^5^U, mcm^5^U, mcm^5^s^2^U, and Um, demonstrating a stronger ability of the updated Dorado model to distinguish Ψ from other uridine modifications. Analysis of Ψ modification probability profiles at non-target sites further supported this improvement: v5.2.0 returned consistently lower probabilities, whereas v5.1.0 produced intermediate values, confirming enhanced specificity in assigning low Ψ probabilities to non-Ψ modifications (Fig. 1C). Nonetheless, many sites that are not assigned a modification (NA U, Figure 1A), showed increased Ψ_ml_ scores, though the values in v5.2.0 were reduced compared to v5.1.0. These Ψ_ml_ scores at NA U sites may result from interference of neighboring modifications.

**Figure 1.**
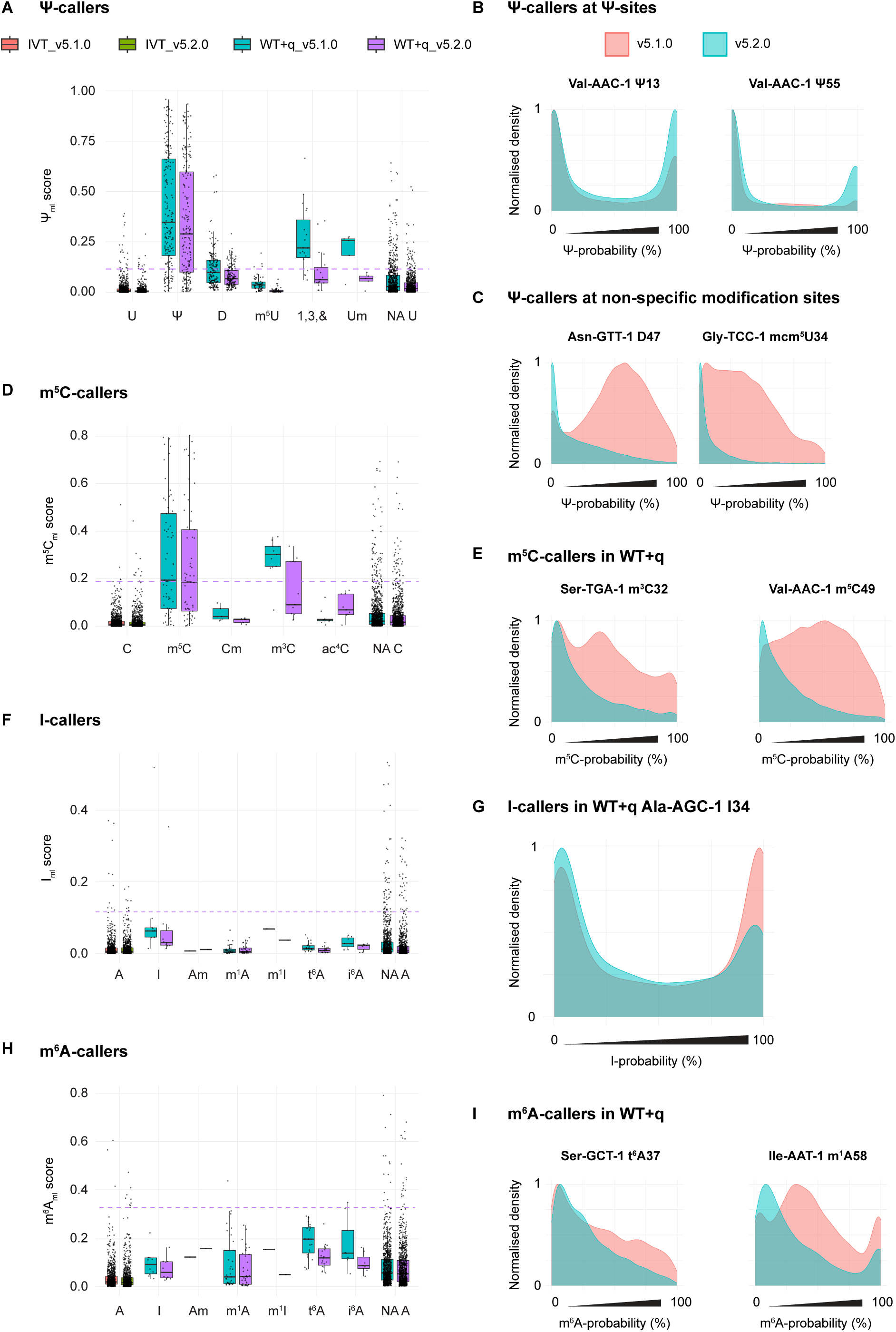
Comparison of the *de novo* modification calling models Dorado v5.1.0 and v5.2.0 on *ex cellulo* tRNA from S*. pombe*. A) Ψ_ml_, scores on *in vitro*-transcribed (IVT) tRNAs and on tRNAs from wild-type *S. pombe* cultured in the presence of queuine (WT + q) at Ψ sites and other annotated tRNA modifications. Sites lacking an annotation are indicated as “NA”. The dashed purple line indicates the ml-score threshold for the Dorado v5.2.0 caller B) Ψ probability distributions at Ψ13 and Ψ55 of tRNA-Val_AAC_ in v5.1.0 (teal) and v5.2.0 (red). C) Ψ probability distributions on dihydrouridine (D47) and mcm^5^U34 using Dorado v5.1.0 and v5.2.0. D) m^5^C_ml_ scores as generated using Dorado v5.1.0 and v5.2.0, represented as in A. E) m^5^C probability distributions on representative m^5^C and m^3^C sites. F) I_ml_ scores as generated using Dorado v5.1.0 and v5.2.0, represented as in A. G) Inosine probability distributions at an I34 site. H) m^6^A_ml_ scores as generated using Dorado v5.1.0 and v5.2.0, represented as in A. I) m^6^A probability distributions at representative t^6^A and m^1^A sites.

In order to call a particular site Ψ-modified or not, we next sought to convert the continuous Ψ_ml_ scores into a binary choice by introducing a Ψ classification threshold. This threshold was defined based on the rate of false-positive *de novo* Ψ predictions on IVT tRNA, which by definition is unmodified (see Methods). Using this threshold for Ψ_ml_ scores, 102 of the 147 annotated Ψ-sites (69.4% true-positive rate) were detected. Furthermore, 27 D sites, two ncm^5^U, one mcm^5^s^2^U site and 33 non-annotated sites yielded scores above the threshold, indicating a substantial false-positive rate of the Ψ caller (Fig. 1 A).

### Evaluation of the Dorado v5.2.0 callers for m^5^C, inosine and m^6^A on tRNA

We next compared the performance of the Dorado v5.2.0 callers for m^5^C, inosine and m^6^A to v5.1.0. Similar to the Ψ caller, ml scores at known modification sites were consistently lower in v5.2.0 than in v5.1.0. For the m^5^C caller, it was interesting to note that ml scores for 3-methyl-cytidine (m^3^C, “m”) were reduced in v5.2.0, whereas those for N^4^-acetylcytidine (ac^4^C, “M”) were increased (Fig. 1 D–E). Also, many non-annotated C sites (NA C) showed increased ml scores, indicating limited specificity of the caller. Using dRNA-seq and m^5^C calling to set a threshold for calling m^5^C, only half of the known m^5^C sites scored above the threshold, indicating a high false-negative rate of the caller. Thus, the v5.2.0 m^5^C caller, as for v5.1.0, shows limited performance in the context of tRNAs.

Similarly, analysis of the inosine caller v5.2.0 indicated limited performance on tRNAs *ex cellulo* (Fig. 1 F). *S. pombe* has seven inosine sites in tRNA, of which only one site (Ala-AGC I34) yielded a high I_ml_ score and surpassed the inosine classification threshold as defined from the analysis of IVT tRNAs (Fig. 1 G).

For evaluation of the m^6^A caller, it is important to consider that tRNAs carry no m^6^A, and any *de novo* calling of this modification by default is an off-target effect. As for the other modification callers, m^6^A_ml_ scores were lower in v5.2.0 compared to v5.1.0 (Fig. 1 H). However, many A sites lacking an annotated modification yielded increased m^6^A_ml_ values. It is furthermore noteworthy that the m^6^A model returned increased scores on the non-modified IVT tRNAs, indicating a high false-positive rate of this caller. Nonetheless, inspection of m^6^A probability distributions at the non-target N^6^-threonylcarbamoyladenosine (t^6^A) and N^1^-methyl-adenosine (m^1^A) modifications showed reduced probability score distributions in the v5.2.0 compared to v5.1.0 model, indicating improved specificity of this caller (Fig. 1 I).

Taken together, these results showed that the v5.2.0 modification calling models yield lower scores than the v5.1.0 models, both at target and off-target modification sites. Altogether, v5.2.0 showed an improved ability to distinguish the target modification from other modifications.

### Evaluation of the Dorado v5.2.0 modification callers for the detection of 2’O-methyl-ribose in tRNA

A major update in Dorado v5.2.0 has been the introduction of modification callers for 2′-O-ribose methylation (Nm). These comprise models for the *de novo* detection of Um, Cm, Am, and Gm sites, which were not available in earlier versions. We therefore evaluated their performance using *ex cellulo* tRNAs from *S. pombe* and IVT tRNAs using the same approach as above.

For the Um-caller, all four annotated Um sites in *S. pombe* tRNA yielded increased Um modification likelihood scores (Um_ml_) that were above the threshold defined by Um_ml_ scores on IVT tRNA (Fig. 2 A). However, multiple Ψ (39 sites), D (23 sites), and m^5^U sites (all except Glu-TTC m^5^U54) also scored above the threshold, revealing a high false-positive rate of this caller. Given the strong response of the Um-caller to m^5^U, this model may therefore be useful for inferring the presence or absence of m^5^U in tRNAs (Fig 2 B).

**Figure 2.**
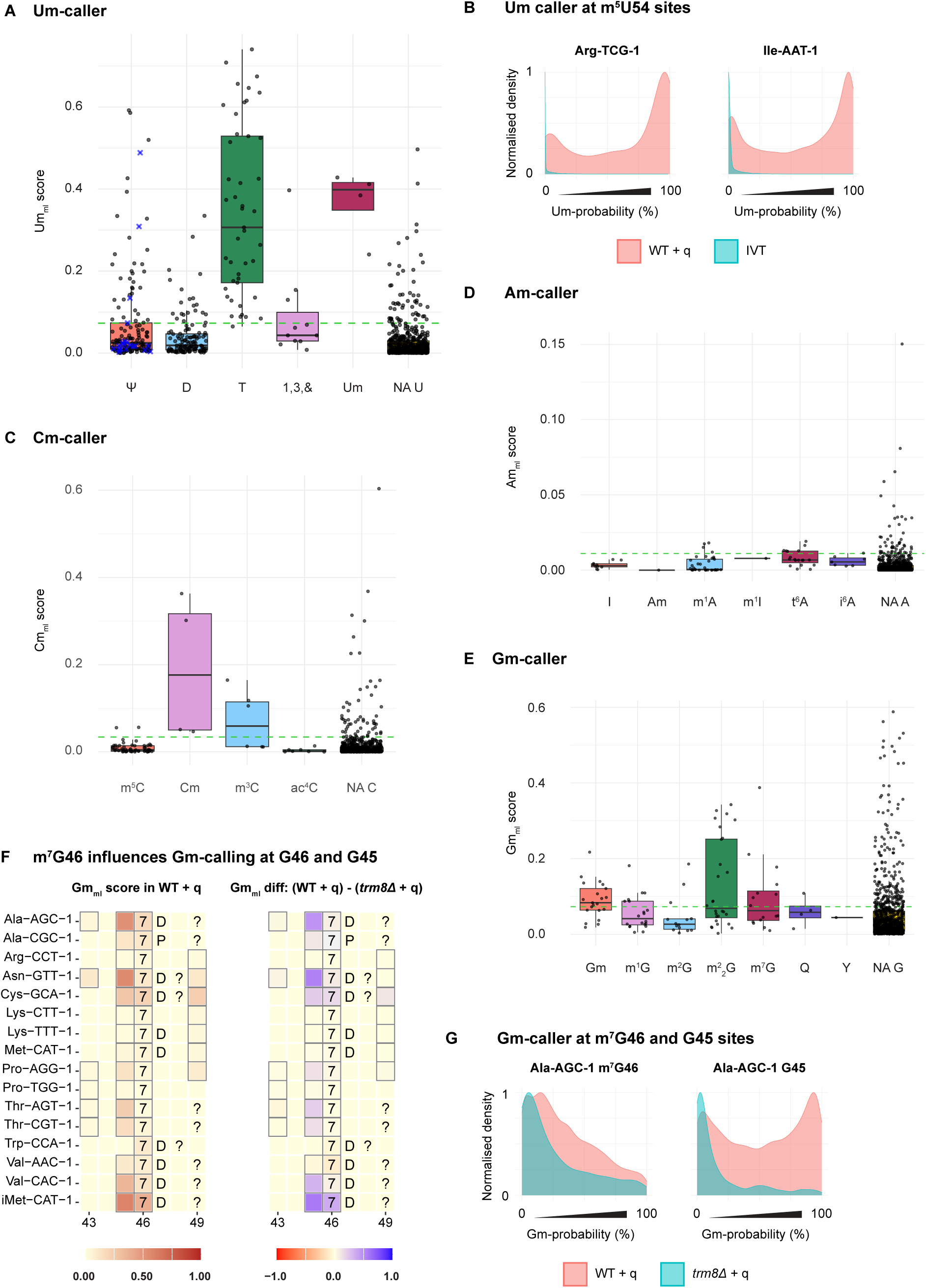
Evaluation of the Dorado v5.2.0 Nm-callers. A) Um_ml_ scores as generated using Dorado v5.1.0 and v5.2.0, represented as in Fig. 1A. B) Um probability distributions at two representative m^5^U sites in tRNAs from wild-type *S. pombe* cultured in the presence of queuine (WT + q) and *in vitro*-transcribed *S. pombe* tRNAs (IVT). C) Cm_ml_ scores, D) Am_ml_ scores, and E) Gm_ml_ scores, as represented in A. F) Heat map of Gm_ml_ scores in WT + q (left) and the difference in Gm_ml_ score between WT + q and *trm8Δ* + q around m^7^G46. Grey boxes indicate sites with low coverage (n ≤ 50). Grey-bordered boxes indicate guanosine sites. G) Comparison of Gm probability distributions in WT + q and *trm8Δ* + q at m^7^G46 and the neighbouring, unmodified G45.

For the *de novo* Cm caller, all four annotated Cm sites in *S. pombe* tRNA displayed increased Cm_ml_ scores that were above the classification threshold, again reflecting good sensitivity (Fig 2 C). The false-positive rate was relatively low, with two m^5^C sites and three m^3^C sites exceeding the threshold.

The *de novo* Am caller exhibited the lowest classification threshold among all modification callers analysed here (0.011), showing a low false positive prediction rate in IVT tRNAs. *S. pombe* tRNAs contain a single annotated Am-site (His-GTG Am5, inferred from *S. cerevisiae* data in Modomics), which scored below this threshold (Fig 2 D). Either the site is misannotated as Am, or the Am caller shows low performance in the context of tRNAs. The scarcity of Am sites in *S. pombe* prevents a more robust evaluation of this caller.

For the *de novo* Gm caller, 15 of the 23 annotated Gm-sites (65.2%) displayed increased Gm_ml_ values that were above the threshold derived from IVT tRNA, reflecting moderate sensitivity (Fig 2 E). Notably, 14 N^2^,N^2^-dimethylguanosine (m^2^_2_G) and 8 N^7^-methylguanosine (m^7^G) positions also gave Gm_ml_ values above the threshold and displayed characteristic Gm modification probability profiles, indicating that the Gm caller erroneously classifies m^2^_2_G and m^7^G as Gm.

Of note, Gm_ml_ scores at G45, which lies adjacent to m^7^G46, were consistently high, suggesting that this resulted from signal spreading from the proximal m^7^G46 (“neighbourhood effect”, see below). To test this, we analysed Gm_ml_ scores from a *trm8Δ S. pombe* strain, which lacks the m^7^G46 modification [31]. Indeed, in *trm8Δ*, Gm_ml_ scores at m^7^G46 were reduced in some tRNAs, and the elevated Gm_ml_ scores at G45 were also reduced in *trm8Δ* compared to wild-type (wt) (Fig 2 F, G). This shows that m^7^G46 interferes with Gm detection both at the modified site itself and at the adjacent G45. Interestingly, not every m^7^G site showed this effect: in some cases (for example, in tRNAs Asn-GTT, Thr-AGT, Thr-CGT, and Val-CAC), high Gm_ml_ scores were observed at G45 without corresponding high Gm_ml_ scores at the neighbouring m^7^G46, and the score was reduced in *trm8Δ*, which lacks m^7^G46 (Fig 2 F). This suggests that the G45–m^7^G46 motif may specifically confuse the Gm-caller in some, but not other tRNAs, leading to false predictions of Gm at position 45 and/or 46.

### Dorado v5.2.0 Ψ-caller is sensitive to neighbouring modifications

In our earlier evaluation of the Dorado v5.1.0 Ψ-caller, we observed that neighbouring modifications such as ncm^5^U34 and Q34 influence *de novo* Ψ-calling at proximal Ψ-sites [20]. We therefore examined whether this effect persists with the updated v5.2.0 models.

The ncm^5^U, mcm^5^U and mcm^5^s^2^U modifications are synthesized by the Elongator complex at the wobble U34 position in selected tRNAs [32–34]. Investigation of the Ψ32 site in tRNAs Leu-TAA, Pro-TGG, and Val-TAC in the wt strain showed that the Ψ caller returned mostly low Ψ probabilities, and that this Ψ site was therefore not detectable in wt. In contrast, in an *elp6Δ* strain, which lacks the Elp6 subunit of the Elongator complex and therefore the U34 modifications [34], the Ψ probability plots showed an increased proportion of reads with high Ψ-scores (Fig 3 A, B). There are two possible interpretations of this observation: either Ψ32 modification increases in the absence of Elongator-dependent U34 modifications, or the U34 modifications interfere with Ψ detection by the Ψ caller. Orthogonal methods are required to distinguish between these possibilities.

**Figure 3.**
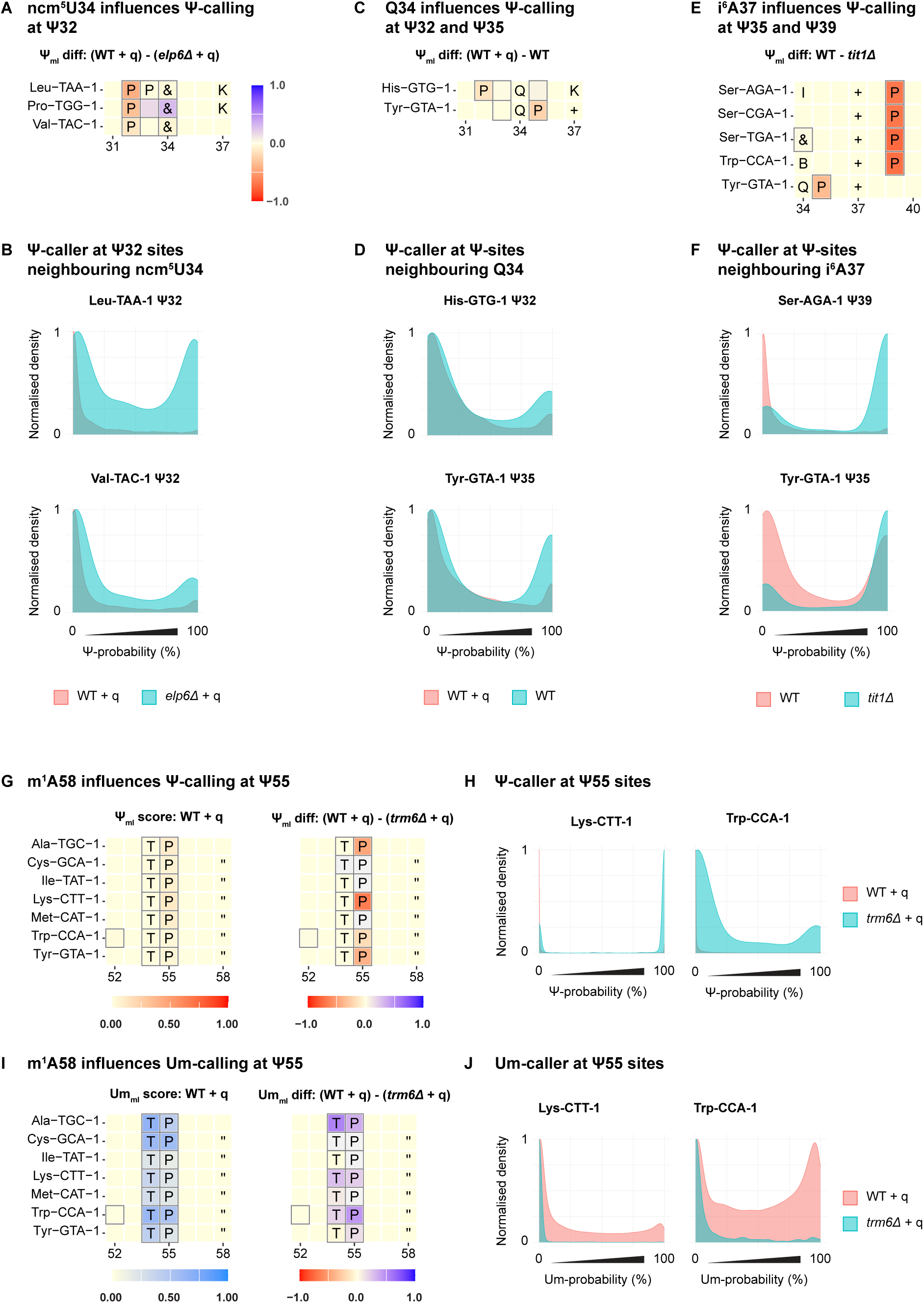
Neighbouring modifications affect Ψ-calling at Ψ sites. A) Heat map visualization of the difference in Ψ_ml_ scores between WT + q and *elp6Δ* + q around ncm^5^U sites (“&”). Grey boxes indicate sites with low coverage (n ≤ 50). Grey-bordered boxes indicate uridine sites in the tRNA. B) Comparison of Ψ probability distributions at Ψ32 in WT + q versus *elp6Δ* + q, which lacks ncm^5^U34. C) Effect of queuosine (Q) modification on Ψ calling at Ψ32 and Ψ35. Representation as in A. D) Ψ probability distributions at Ψ32 and Ψ35 in the presence (WT + q) and absence (WT) of Q34 modification. E) Effect of i^6^A modification on Ψ calling at Ψ35 and Ψ39. Representation as in A, comparing wild-type *S. pombe* tRNAs (WT) to tRNA lacking i^6^A (*tit1Δ*). F) Ψ probability distributions at Ψ35 and Ψ39 in the presence (WT) and absence (*tit1Δ*) of i^6^A37. G) Effect of m^1^A58 (“) on Ψ calling at Ψ55. Left, Ψ_ml_ scores around Ψ55 in selected tRNAs. Representation as in A. Right, difference of Ψ_ml_ scores between WT + q and *trm6Δ* + q, which lack m^1^A58. H) Comparison of Ψ probability distributions at Ψ55 sites in the presence (WT + q) or absence (*trm6Δ* + q) of m^1^A58. I) Erroneous Um calling at Ψ55 is affected by m^1^A58. Representation as in G. J) Um probability distribution at selected Ψ55 in the presence (WT + q) or absence (*trm6Δ* + q) of m^1^A58.

Similarly, in the absence of queuosine modification at position 34 (Q34), which was achieved by culturing *S. pombe* cells without queuine [30], Ψ probability plots revealed higher Ψ probabilities at neighbouring Ψ32 and Ψ35 in wt without compared to with Q34 modification (Fig 3 C, D). As above, this indicates either that Q34 interferes with Ψ calling at neighbouring positions, or Ψ levels are increased at these sites in the absence of Q34.

Furthermore, Ψ_ml_ scores were consistently low at Ψ35 and Ψ39 sites located proximal to N^6^-isopentenyladenosine at position 37 (i^6^A37). In a *tit1Δ* strain, which lacks i^6^A37 [35], Ψ_ml_ scores at these neighbouring Ψ35 and Ψ39 sites were elevated relative to wt (Fig 3 E, F). Again, whether these effects reflect genuine increases in Ψ levels in the absence of i^6^A37, or rather interference of neighbouring modifications with Ψ detection cannot be resolved in this analysis.

Surprisingly, we also detected low Ψ_ml_ scores at several Ψ55 sites, with the values in six tRNAs (Ala-TGC, Cys-GCA, Ile-TAT, Lys-TT, Met-CAT, Trp-CCA, and Tyr-GTA) falling below the threshold defined by IVT tRNAs (Fig. 3G, H). These reductions were consistently accompanied by elevated Um_ml_ scores at the same positions, suggesting misclassification of Ψ55 as Um55 (Fig. 3I, J). Given that Ψ55 is a highly conserved site of pseudouridylation [36–38], it is unlikely that these sites are Um-modified. Based on earlier observations, we hypothesised that the presence of proximal m^1^A58 might underlie this effect. Indeed, in a *trm6Δ* strain, which lacks m^1^A58 [39], Ψ_ml_ scores at Ψ55 sites with low values in wt showed increased Ψ_ml_ scores, while the corresponding Um_ml_ scores decreased in *trm6Δ* relative to wt (Fig. 3 H, J). These results indicate that m^1^A58 interferes with Ψ55 detection in some tRNAs and causes the Dorado Ψ-caller to misclassify Ψ55 as Um55. An alternative interpretation is that m^1^A58 is inhibitory for Ψ55 formation in a subset of tRNAs.

## Discussion

The detection of RNA modifications in biological samples is critical for elucidating their contributions to gene regulation and disease. Direct RNA sequencing using biological nanopores, currently implemented by Oxford Nanopore Technologies, provides a powerful method to identify such modifications. Here, we systematically assessed the performance of the latest basecalling models (Dorado v5.2.0) for the *de novo* detection of eight RNA modifications (Ψ, m^5^C, I, m^6^A, Um, Am, Cm, Gm), using *Schizosaccharomyces pombe* tRNAs as a model system.

Among the v5.2.0 models, the Ψ caller performed most reliably, exhibiting high sensitivity toward genuine Ψ modifications and minimal erroneous detection at many other uridine sites. In particular, compared to v5.1.0, the Ψ caller has an improved ability to distinguish between Ψ and Um, as well as the Elongator-dependent U34 modifications. The distinction of Ψ from Um most likely results from the fact that a separate model was trained to detect Um. Notably, dihydrouridine (D) positions still produced elevated Ψ scores, and the modification probability distributions resemble those observed at Ψ sites with low modification levels. This suggests that the Dorado v5.2.0 model may also be employed to detect D modifications, although the relationship between signal intensity and D modification level remains to be determined. However, though improved compared to the v5.1.0 model, the v5.2.0 Ψ caller remains sensitive to modifications on other nucleobases in the vicinity.

Therefore, confidently assigning a Ψ site ideally still requires combining *de novo* and comparative methods by sequencing (t)RNAs from wild-type samples and samples lacking Ψ. Bulky modifications such as Q and i^6^A continued to interfere with Ψ detection, and new cross-effects were observed – for instance, the possible misclassification of Ψ55 as Um55 in the presence of m^1^A58 in a subset of tRNAs. These findings highlight the importance of considering modification sequence context when interpreting modification predictions.

In contrast to the v5.2.0 Ψ caller, the m^5^C and inosine callers showed limited performance on tRNAs, as was seen with v5.1.0, with modification likelihoods failing to exceed the false positive rate for many cognate sites. Furthermore, the m^6^A caller exhibited a high false positive rate on non-modified sequences, thus limiting its use. These shortcomings emphasise the importance of validating Dorado modification models on biological samples before using them for *de novo* modification prediction.

Together, our results underline both the potential and the current limitations of the Dorado v5.2.0 models. While certain callers – particularly the Ψ-caller – show good performance for identifying modification sites *de novo*, direct modification basecalling should be interpreted with caution and ideally requires validation using orthogonal methods to ensure confident assignment of RNA modifications.

## Availability of data and materials

The sequencing data is available at the European Nucleotide Archive (accession number PRJEB104115). The analysis scripts are available at https://github.com/bhargesh-patel/Dorado-v5.2.0-evaluation.

## Funding

This work was funded by Deutsche Forschungsgemeinschaft (grants EH237/19-1 and EH237/21-1 to A.E.E.-M.), Deutscher Akademischer Austauschdienst DAAD to B.I.P., and the Chinese Scholarship Council to Y.S..

## Authors’ contributions

A.E.E.-M. conceived the project. F.N.M.R., Y.S. and B.I.P. performed the sequencing of samples. B.I.P. implemented the analysis pipeline. A.E.E.-M. and B.I.P. drafted the manuscript with contributions from all authors.

## Acknowledgements

We thank the high-performance computing facility of Humboldt-Universität zu Berlin for computational resources.

## Notes

https://github.com/bhargesh-patel/Dorado-v5.2.0-evaluation

### Competing Interest Statement

The authors have declared no competing interest.

## References

1. Sloan KE, Warda AS, Sharma S, Entian K-D, Lafontaine DLJ, Bohnsack MT. Tuning the ribosome: The influence of rRNA modification on eukaryotic ribosome biogenesis and function. RNA Biol. 2017;14:1138–52. 10.1080/15476286.2016.1259781.

2. Zhang M, Lu Z. tRNA modifications: greasing the wheels of translation and beyond. RNA Biol. 2025;22:1–25. 10.1080/15476286.2024.2442856.

3. Suzuki T. The expanding world of tRNA modifications and their disease relevance. Nat Rev Mol Cell Biol. 2021;22:375–92. 10.1038/s41580-021-00342-0.

4. Gilbert W V., Nachtergaele S. mRNA Regulation by RNA Modifications. Annu Rev Biochem. 2023;92:175–98. 10.1146/annurev-biochem-052521-035949.

5. Roundtree IA, Evans ME, Pan T, He C. Dynamic RNA Modifications in Gene Expression Regulation. Cell. 2017;169:1187–200. 10.1016/j.cell.2017.05.045.

6. Motorin Y, Helm M. tRNA Stabilization by Modified Nucleotides. Biochemistry. 2010;49:4934–44. 10.1021/bi100408z.

7. Qiu L, Jing Q, Li Y, Han J. RNA modification: mechanisms and therapeutic targets. Molecular biomedicine. 2023;4:25. 10.1186/s43556-023-00139-x.

8. Delaunay S, Pascual G, Feng B, Klann K, Behm M, Hotz-Wagenblatt A, et al. Mitochondrial RNA modifications shape metabolic plasticity in metastasis. Nature. 2022;607:593–603. 10.1038/s41586-022-04898-5.

9. Chujo T, Tomizawa K. Human transfer RNA modopathies: diseases caused by aberrations in transfer RNA modifications. FEBS J. 2021;288:7096–122. 10.1111/febs.15736.

10. Lv X, Zhang R, Li S, Jin X. tRNA Modifications and Dysregulation: Implications for Brain Diseases. Brain Sci. 2024;14:633. 10.3390/brainsci14070633.

11. Deamer D, Akeson M, Branton D. Three decades of nanopore sequencing. Nat Biotechnol. 2016;34:518–24. 10.1038/nbt.3423.

12. van Dijk EL, Jaszczyszyn Y, Naquin D, Thermes C. The Third Revolution in Sequencing Technology. Trends in Genetics. 2018;34:666–81. 10.1016/j.tig.2018.05.008.

13. Wang Y, Zhao Y, Bollas A, Wang Y, Au KF. Nanopore sequencing technology, bioinformatics and applications. Nat Biotechnol. 2021;39:1348–65. 10.1038/s41587-021-01108-x.

14. Jain M, Olsen HE, Paten B, Akeson M. The Oxford Nanopore MinION: delivery of nanopore sequencing to the genomics community. Genome Biol. 2016;17:239. 10.1186/s13059-016-1103-0.

15. Rang FJ, Kloosterman WP, de Ridder J. From squiggle to basepair: computational approaches for improving nanopore sequencing read accuracy. Genome Biol. 2018;19:90. 10.1186/s13059-018-1462-9.

16. Loman NJ, Quick J, Simpson JT. A complete bacterial genome assembled de novo using only nanopore sequencing data. Nature Methods 2015 12:8. 2015;12:733–5. 10.1038/nmeth.3444.

17. Leger A, Amaral PP, Pandolfini L, Capitanchik C, Capraro F, Miano V, et al. RNA modifications detection by comparative Nanopore direct RNA sequencing. Nat Commun. 2021;12:7198. 10.1038/s41467-021-27393-3.

18. Kovaka S, Hook PW, Jenike KM, Shivakumar V, Morina LB, Razaghi R, et al. Uncalled4 improves nanopore DNA and RNA modification detection via fast and accurate signal alignment. Nat Methods. 2025;22:681–91. 10.1038/s41592-025-02631-4.

19. Piechotta M, Naarmann-de Vries IS, Wang Q, Altmüller J, Dieterich C. RNA modification mapping with JACUSA2. Genome Biol. 2022;23:115. 10.1186/s13059-022-02676-0.

20. Rübsam FNM, Liu-Wei W, Sun Y, Patel BI, van der Toorn W, Piechotta M, et al. MoDorado: enhanced detection of tRNA modifications in nanopore sequencing by off-label use of modification callers. Nucleic Acids Res. 2025;53. 10.1093/nar/gkaf795.

21. Pan T. Modifications and functional genomics of human transfer RNA. Cell Res. 2018;28:395–404. 10.1038/S41422-018-0013-Y;SUBJMETA=1793,337,384,521,574,631;KWRD=SMALL+RNAS,TRNAS.

22. Esfahani NG, Stein AJ, Akeson S, Tzadikario T, Jain M. Evaluation of Nanopore direct RNA sequencing updates for modification detection. bioRxiv. 2025. 10.1101/2025.05.01.651717.

23. Zou Y, Ahsan MU, Chan J, Meng W, Gao S-J, Huang Y, et al. A comparative evaluation of computational models for RNA modification detection using nanopore sequencing with RNA004 chemistry. Brief Bioinform. 2025;26. 10.1093/bib/bbaf404.

24. Chan PP, Lowe TM. GtRNAdb 2.0: an expanded database of transfer RNA genes identified in complete and draft genomes. Nucleic Acids Res. 2016;44:D184–9. 10.1093/nar/gkv1309.

25. Rutherford KM, Lera-Ramírez M, Wood V. PomBase: a Global Core Biodata Resource—growth, collaboration, and sustainability. Genetics. 2024;227. 10.1093/genetics/iyae007.

26. Daily J. Parasail: SIMD C library for global, semi-global, and local pairwise sequence alignments. BMC Bioinformatics. 2016;17:81. 10.1186/s12859-016-0930-z.

27. Sun Y, Piechotta M, Naarmann-de Vries I, Dieterich C, Ehrenhofer-Murray AE. Detection of queuosine and queuosine precursors in tRNAs by direct RNA sequencing. Nucleic Acids Res. 2023;51:11197–212. 10.1093/nar/gkad826.

28. Li H. Aligning sequence reads, clone sequences and assembly contigs with BWA-MEM. 2013.

29. Lucas MC, Pryszcz LP, Medina R, Milenkovic I, Camacho N, Marchand V, et al. Quantitative analysis of tRNA abundance and modifications by nanopore RNA sequencing. Nat Biotechnol. 2023;42:72–86. 10.1038/s41587-023-01743-6.

30. Müller M, Hartmann M, Schuster I, Bender S, Thüring KL, Helm M, et al. Dynamic modulation of Dnmt2-dependent tRNA methylation by the micronutrient queuine. Nucleic Acids Res. 2015;43:10952–62. 10.1093/nar/gkv980.

31. de Zoysa T, Phizicky EM. Hypomodified tRNA in evolutionarily distant yeasts can trigger rapid tRNA decay to activate the general amino acid control response, but with different consequences. PLoS Genet. 2020;16:e1008893. 10.1371/JOURNAL.PGEN.1008893.

32. Gaik M, Kojic M, Wainwright BJ, Glatt S. Elongator and the role of its subcomplexes in human diseases. EMBO Mol Med. 2023;15. 10.15252/EMMM.202216418/ASSET/1B037674-8E48-420A-BEDB-9DF6A69860DA/ASSETS/GRAPHIC/EMMM202216418-FIG-0001-M.PNG.

33. Huang B, Johansson MJO, Byström AS. An early step in wobble uridine tRNA modification requires the Elongator complex. RNA. 2005;11:424–36. 10.1261/rna.7247705.

34. Bauer F, Matsuyama A, Candiracci J, Dieu M, Scheliga J, Wolf DA, et al. Translational Control of Cell Division by Elongator. Cell Rep. 2012;1:424–33. 10.1016/j.celrep.2012.04.001.

35. Lamichhane TN, Blewett NH, Crawford AK, Cherkasova VA, Iben JR, Begley TJ, et al. Lack of tRNA Modification Isopentenyl-A37 Alters mRNA Decoding and Causes Metabolic Deficiencies in Fission Yeast. Mol Cell Biol. 2013;33:2918–29. 10.1128/MCB.00278-13;SUBPAGE:STRING:ACCESS.

36. Roovers M, Droogmans L, Grosjean H. Post-Transcriptional Modifications of Conserved Nucleotides in the T-Loop of tRNA: A Tale of Functional Convergent Evolution. Genes (Basel). 2021;12:140. 10.3390/GENES12020140.

37. Mukhopadhyay S, Deogharia M, Gupta R. Mammalian nuclear TRUB1, mitochondrial TRUB2, and cytoplasmic PUS10 produce conserved pseudouridine 55 in different sets of tRNA. RNA. 2021;27:66–79. 10.1261/rna.076810.120.

38. Becker HF, Motorin Y, Planta RJ, Grosjean H. The yeast gene YNL292w encodes a pseudouridine synthase (Pus4) catalyzing the formation of Ψ 55 in both mitochondrial and cytoplasmic tRNAs. Nucleic Acids Res. 1997;25:4493–9. 10.1093/NAR/25.22.4493.

39. Tasak M, Phizicky EM. Initiator tRNA lacking 1-methyladenosine is targeted by the rapid tRNA decay pathway in evolutionarily distant yeast species. PLoS Genet. 2022;18:e1010215. 10.1371/JOURNAL.PGEN.1010215.

